# Opioid-blunted cortisol response to stress is associated to increased negative mood and wanting of social reward

**DOI:** 10.1101/2021.08.13.456110

**Authors:** Claudia Massaccesi, Matthaeus Willeit, Boris B. Quednow, Urs M. Nater, Claus Lamm, Daniel Mueller, Giorgia Silani

**Author notes:** Correspondence: Claudia Massaccesi Giorgia Silani.

## Abstract

Animal research suggests a central role of the μ-opioid receptor (MOR) system in regulating affiliative behaviors and in mediating the stress-buffering function of social contact. However, the neurochemistry of stress-related social contact seeking in humans is still poorly understood. In a randomized, double-blind, between-subject design, healthy female volunteers (N = 80) received either 10 mg of the μ-opioid agonist morphine sulfate, or a placebo. Following a standardized psychosocial stress induction, participants engaged in a social reward task, in which the motivation to obtain skin-to-skin social touch and the hedonic reactions elicited by such touch were assessed.

Morphine prevented the increase of salivary cortisol typically observed following acute stress exposure. Notably, this altered HPA axis responsivity was associated with increased negative affect in response to psychosocial stress, and with enhanced subjective wanting of highly rewarding social contact.

These findings provide novel evidence on the effect of exogenous opioids administration on the reactions to psychosocial stress and point to a state-dependent regulation of social motivation.

## Introduction

Social behaviors such as bonding and affiliation are crucial for the survival and wellbeing of many species. By providing fundamental benefits (e.g., promoting safety and enhancing stress resilience) and by generating comfort and pleasure, social stimuli (e.g., social contact) gain rewarding value, inducing approach motivation. Inability to form and maintain social bonds contributes to a range of psychiatric and physical pathologies [e.g., 1,2], highlighting the importance to better understand the neurobiological basis of social motivation.

Based on pharmacological evidence in isolated animals showing that exogenous μ-opioids administration (e.g. morphine) reduces separation distress and contact seeking, the Brain Opioid Theory of Social Attachment [3] pinpoints the μ-opioid receptor (MOR) system as a key mediator of bonding and affiliation. Extending this model, Løseth and colleagues propose that the MOR system regulates social behaviors in a manner that is context-dependent [4]. Specifically, in contexts of distress (such as the social isolation described above), social stimuli are sought because they stimulate endogenous μ-opioids release, which in turn reduces pain and negative affect. If opioids are exogenously provided, this will result in alleviated distress, and therefore reduced need for social contact. On the other hand, in contexts of comfort, endogenous MOR activity mediates the rewarding properties, and associated pleasure, of social stimuli. In this case, exogenous MOR stimulation will result in increased pleasure and motivation to seek for social contact.

In the last decade, preliminary confirmatory evidence on the state-dependent MOR regulation of affiliation and social reward processing in humans has been provided. During states of comfort, MOR blockade decreases wanting and/or liking of different social stimuli [5–7, but see also 8], as well as feelings of social connection [9–11], interpersonal closeness and social reward expectation [12], while MOR enhancement increases wanting and liking of attractive faces [6]. Using PET, Hsu and colleagues [13] showed that endogenous MOR activity during social rejection is positively associated with reduced negative affect, while during social acceptance it predicts greater desire for social interaction. However, to date, causal evidence of MOR regulation of social motivation and contact seeking during distress is lacking.

Here, we aimed at filling this knowledge gap by investigating the effect of MOR agonist administration (morphine) on social motivation and social pleasure following stress exposure. Using a double-blind, placebo-controlled, randomized, between-subjects design, female participants (N = 80) were orally administered with either 10 mg morphine sulfate (a highly selective μ-opioid agonist) or placebo. Following a psychosocial stress induction procedure, the motivation to obtain social touch (wanting) and the pleasure elicited by receiving it (liking) were assessed (see Figure 1 for a detailed description). To enhance the comparability with animal research, i) a stressor of social nature was employed; ii) in addition to self-reports of wanting and liking, we assessed real physical effort and hedonic facial reactions, approximating the motivational and hedonic (i.e., wanting and liking) measures used in animal studies; iii) to parallel grooming in animals, skin-to-skin touch was employed as a social reward.

**Figure 1.**
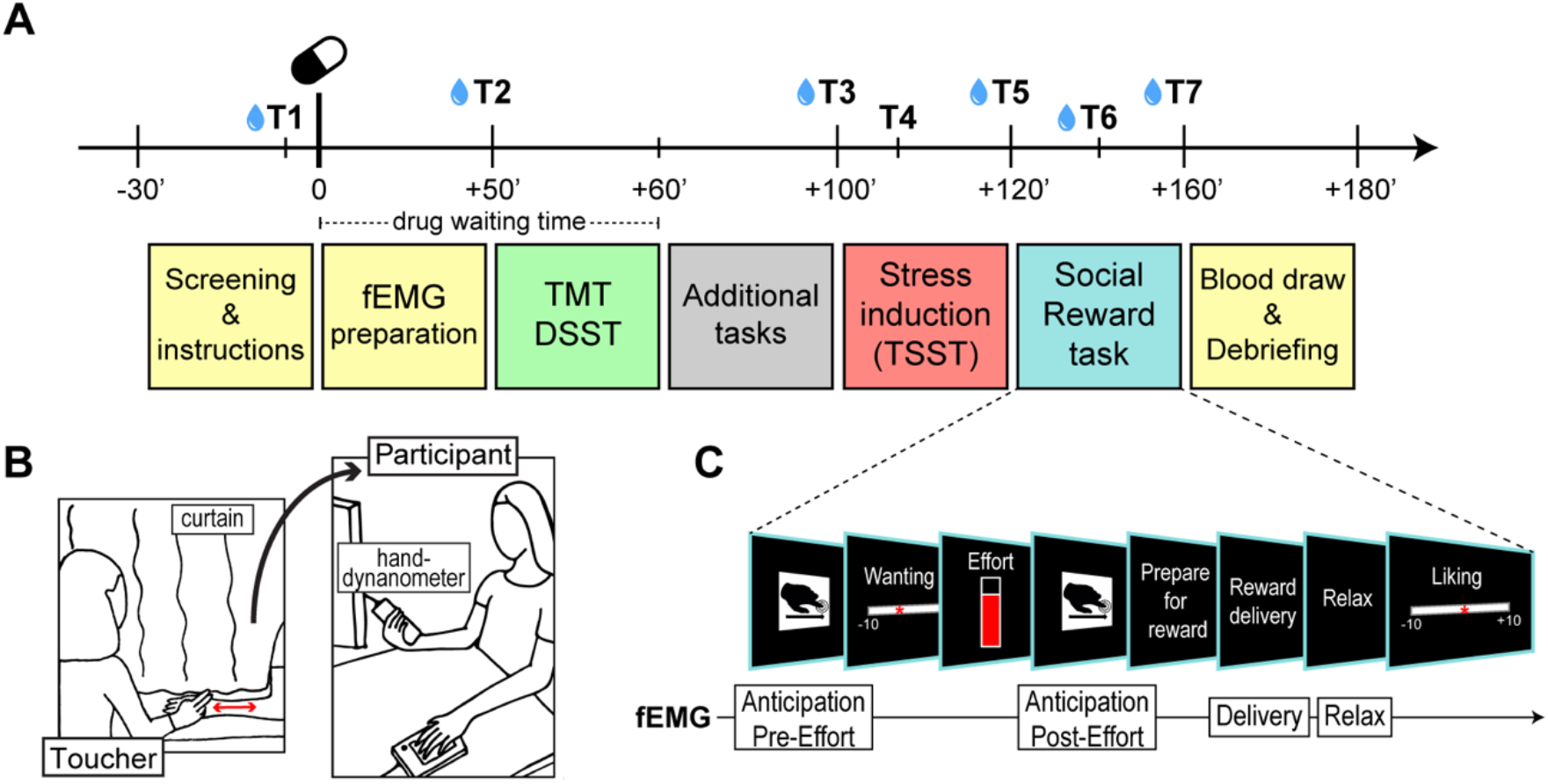
Overview of the experimental session, set-up, and trial structure of the Social Reward task. (A) Overview of the experimental session. T1 – T7 represent the time points at which subjective and/or physiological measures were obtained. Blue drops indicate saliva sample collection. (B) Set-up of the Social Reward task. Participants were seated in front of the monitor, holding the hand-dynamometer in the right hand. The left arm was rested on a cushion, next to a keyboard used to express judgements during the task. The toucher was seated on the other side of a curtain used to limit the participants’ field of view to the monitor. Touch was administered on the participants’ left forearm using the index and middle finger, at 3 stroking speeds corresponding to 3 levels of reward (high = 6 cm/s, low = 21 cm/s, very low = 27 cm/s). (C) Trial structure of the Social Reward task. Facial electromyography (fEMG) was recorded during the whole task and analyzed in reward anticipation (Anticipation Pre-Effort and Post-Effort) and consumption (Delivery and Relax). TMT, Trail Making Test; DSST, Digit Symbol Substitution Test; TSST, Trier Social Stress Test.

As recent human evidence showed that exogenous MOR stimulation alleviates stress, as indicated by decreased cortisol response and threat/challenge appraisal [14,15], we expected reduced responses to stress after morphine administration, compared to placebo. Based on previous theoretical models and animal evidence [3,4], we hypothesized that this opioid-blunted stress response would be associated to decreased social motivation. Given that previous studies showed an effect of stress on wanting rather than on liking of touch [16] or food [e.g. 17] reward, we did not expect changes in the hedonic reactions to social contact. Finally, given that MOR manipulations have been shown to induce the strongest effects on the best reward option available [6,18], and considering the stress buffering function of C tactile (CT)-optimal touch [19], we expected the predicted effects to be stronger for the most valuable social reward (i.e., touch at 6 cm/s, which is in the CT-optimal range stroking speed of 1 to 10 cm/s).

## Materials and Methods

### Study design

The between-subject, double-blind, placebo-controlled study consisted of one experimental session in which participants received either 10 mg morphine sulfate or a placebo.

### Participants

Based on previous work that had investigated the effects of stress on social reward processing [16] and the effects of similar compounds on stress responses [14], we aimed at collecting data from 40 participants per group. The study sample included 82 participants, of which 42 received morphine (MORPH) and 40 received a placebo (PLB). Two participants (MORPH) did not complete the session and were therefore not included in data analysis, yielding to a final sample size of 80 participants (40/group)^1^. Only female participants were included due to i) gender differences in opioid pharmacokinetics [20,21] and stress response [22], ii) expected higher preference of same-gender touch in females than males [23,24]. Participants were tested during the luteal phase of their menstrual cycle, as determined by self-report of their last menstruation and average cycle length. All participants reported to be right-handed, to smoke less than ten cigarettes per week, to have no history of current or former drug abuse, to have a BMI between 17 and 35, and to be free of psychiatric or neurological disorders. Other exclusion criteria were: single or repeated use of any strong opioids in the last two years, use of hormonal contraceptives, regular intake of medications, current pregnancy or breastfeeding, suffering from impaired respiratory functions, respiratory weakness or lung disease, injury/disease of the arms (making it impossible to squeeze with the right hand and to be caressed on the left forearm). Participants were instructed to refrain from eating, brushing their teeth and consuming caffeinated beverages, juices, and chewing gum in the two hours preceding the test, as well as from smoking, doing physical activity and intaking alcohol and medications in the 24 hours preceding the test. The two experimental groups did not differ significantly in terms of age, BMI, autistic traits (short version of the German Autism Spectrum Quotient, AQ-k) [25], general (State-Trate Anxiety Inventory, STAI) [26] and social (Liebowitz Social Anxiety Scale, LSAS) [27] anxiety, and social touch appreciation (Social Touch Questionnaire, STQ) [28] (see Table 1). The study was approved by the Ethics Committee of the Medical University of Vienna (EK 1393/2017) and was performed in line with the Declaration of Helsinki [29]. All participants signed a consent form before taking part in the study.

**Table 1.** Demographic and self-reported substance use characteristics of the participants.

|  | MORPHINE | PLACEBO | *p* value |
| --- | --- | --- | --- |
| N | 40 | 40 | --- |
| Age (years) | 23.1 ± 3.1 | 23.9 ± 3.9 | 0.31 |
| BMI (kg/m<sup>2</sup>) | 21.4 ± 3.4 | 22.3 ± 2.7 | 0.19 |
| Autism (AQ-k) | 6.5 ± 3.8 | 5.7 ± 3.6 | 0.34 |
| Anxiety (STAI) | 38.1 ± 9.4 | 35.7 ± 8.3 | 0.23 |
| Social Anxiety (LSAS) | 36.7 ± 17.9 | 33.2 ± 19.0 | 0.39 |
| Social Touch Preferences (STQ) | 25.3 ± 9.5 | 27.1 ± 7.7 | 0.36 |
| Drug use (% lifetime – last year) |  |  |  |
| Cannabis | 47.5 – 25 | 60 – 32.5 | --- |
| Tranquilizers | 7.5 – 5 | 5 – 2.5 | --- |
| Stimulants | 25 – 10 | 17.5 – 7.5 | --- |
| Opiates | 7.5 – 0 | 2.5 – 0 | --- |
| Hallucinogens | 12.5 – 2.5 | 12.5 – 2.5 | --- |
| Other | 5 – 2.5 | 2.5 – 0 | --- |

### Procedure

The study was conducted at the Department of Psychiatry and Psychotherapy of the Medical University of Vienna. After completing an online survey to assess their eligibility, potential participants were first invited to a health screening (~45’), including blood examination, electrocardiogram, blood pressure measurement, and psychiatric interview (Mini-International Neuropsychiatric Interview) [30]. Eligible participants were then invited to the experimental session (~210’), which always started between 11:30 and 12:30 in order to control for cortisol diurnal fluctuations [31]. At the beginning of the session, urine drug and pregnancy tests were administered. After baseline mood and physiological measures were collected, participants received a standardized snack, and took the assigned capsule. Throughout the session, mood and physiological measures were obtained at regular intervals. The experimental tasks were completed between 60 and 160 min after drug administration, and included economic decision making, facial mimicry, emotion recognition, stress induction and social reward processing (see Figure 1A for an overview of the session timeline). Here we will focus on the last two, while the others will be reported elsewhere. Approximately 180 min after pill administration, a blood sample was taken to confirm drug uptake (see Table S2 of the *Supplementary Material*). After completing the experimental session, participants were debriefed and received a financial compensation of 65€. Half of the sample was tested before and half during the COVID-19 pandemic (see *Supplementary Material* for details regarding the employed safety measures and additional statistical analyses to assess possible effects of the pandemic and safety measures on the results).

### Drug administration

Ten mg of morphine sulfate (Morapid®) were orally administered, with the specific dosage chosen so as to stimulate the activity of the MOR system with minimal subjective (side-)effects. Morphine is a selective MOR agonist and, for oral administration, has an average bioavailability of 30–40%, a maximal effect (t-max) at 1–2 h after administration, and a half-life of 2–4 h [32]. Placebo consisted of capsules containing 650 mg of mannitol (sugar), visually identical to the ones containing morphine.

### Stress induction

In order to induce a stress reaction in the participants, the Trier Social Stress Test (TSST) [33] was employed. In the TSST, participants were given 3 minutes to prepare a 5 min speech for a mock-job interview, followed by a 5 min arithmetic task, in which they were asked to count backwards from 2043 in steps of 17 as fast and as accurate as possible. The speech and arithmetic tasks were completed in front of an evaluating panel (one male and one female confederate). Participants were told that these tasks would be video recorded via a camera located next to the examiners (no video was actually taped).

### Social Reward task

#### Stimuli

Pleasantness of touch is highest between 1 and 10 cm/s, and decreases between 10 and 30 cm/s [34]. Gentle caresses in three different speeds (CT-optimal: 6 cm/s, non-CT-optimal: 21 cm/s and 27 cm/s) were thus used as social rewards of different levels of pleasantness (high, low, very low). The suitability of these stroking speeds has been confirmed in three previous independent studies from our group [7,16,35]. Caresses were delivered over a previously-marked (from the wrist towards the elbow) 9 cm area on the participant’s left forearm by a female experimenter, moving her index and middle fingers back and forth in the marked area (Figure 1B). Touch delivery was guided by auditory rhythms, which matched the frequency of the stimulation, over headphones. The experimenter administering the touch was seated on the other side of a curtain, used to limit the participant’s field of view to the monitor (Figure 1B). All experimenters were presented as trained masseurs, wore standardized clothes (white t-shirt and beige trousers) to minimize differences in their appearance, and underwent extensive training on the tactile stimulation delivery.

#### Task

The Social Reward task [7,16,35] consisted of two blocks of 16 trials, separated by a 5 min break. To avoid habituation to the touch, the site of application (left or right area of the forearm) was alternated within the two blocks, in a counter-balanced order. Before starting the task, participants experienced each type of touch once and performed two training trials. Each trial was structured as follow (see Figure 1C): i) announcement of the best attainable reward (high or low, 16 trials each, 3 s); ii) rating of subjective wanting via a VAS ranging from −10 (not at all) to +10 (very much) (no time limit); iii) effort task (4 s), requiring to squeeze a hand-dynamometer (HD-BTA, Vernier Software & Technology, USA) with the right hand, in order to obtain the announced reward – the applied force, displayed via an online visual-feedback, was expressed as percentage of the participants’ maximum voluntary contraction (MVC), measured immediately before the task, and translated into the probability of obtaining the announced reward (0–100%); iv) announcement of the reward obtained (high, low, or – if insufficient force had been exerted – very low; 2 s); v) reward delivery (6 s); vi) relax phase (5 s); vii) rating of subjective liking via a VAS ranging from −10 (not at all) to +10 (very much) (no time limit). At the end of the task, participants’ MVC was measured again.

The task was implemented in Matlab 2014b (MathWorks, Inc) and presented on an LCD monitor with a resolution of 1280 × 1024 pixels.

### Facial electromyography (EMG)

Facial EMG was recorded throughout the Social Reward task, using a g.USBamp amplifier (g.tec Medical Engineering GmbH) and the software Matlab (MathWorks, Inc). Participant’s face areas were prepared using alcohol, water and an abrasive paste. Reusable Ag/AgCl electrodes were then attached bipolarly according to guidelines [36] on the left corrugator supercilii and zygomaticus major muscles. A ground electrode was attached to the participant’s forehead and a reference electrode on the left mastoid. The EMG data were sampled at 1200 Hz with impedances below 20 kΩ. Data preprocessing included filtering with a 20 to 400 Hz bandpass filter, rectification and smoothing with a 40 Hz low-pass filter.

Each trial was divided in 4 epochs: Anticipation Pre-Effort (announcement of best attainable reward, 3 s), Anticipation Post-Effort (announcement of attained reward, 2 s), Delivery (touch administration, 6 s), and Relax (relax phase after reward delivery, 5 s). EMG was first averaged over 1 s time-windows and then over the epoch total duration. For each trial, values in the four epochs were expressed as percentage of the average amplitude during the fixation cross at the beginning of that trial (baseline, 2 s). Outliers in baseline values (defined as values more than 3 SDs away from the subjects’ average baseline) were substituted with the average amplitude of the baseline preceding and following that trial. The extracted epochs were visually inspected to identify signal artifacts which were then removed (33 epochs for corrugator and 51 epochs for zygomaticus across 5 participants). Because of data skewness and to reduce the effect of non-experimental movements, for each subject, epochs over the subject’s mean ± 3 SD were removed (average removed epochs per subject: corrugator: *M* = 2.4, *SD* = 1.2; zygomaticus: *M* = 3.1, *SD* = 1.1), and the remaining data were transformed using natural logarithmic transformation.

### Subjective measures of stress and mood

During the session, participants completed self-report questionnaires assessing their mood and subjective state at regular intervals. Happiness, calmness, relaxation, feeling good, as well as stress, tension, anxiety and feeling bad were assessed using VAS ranging from “not at all” (+1) to “very much” (+101) at 7 time points (T1-T7; see Figure 1A). Positive and negative mood items were averaged to constitute the “Positive mood” and “Negative mood” scales used for statistical analyses. The German short version Profile of Mood States (POMS) [37], consisting of 4 subscales for current mood (depression, vigor, fatigue, and displeasure), was administered at 5 time points (T1, T2, T3, T5, T6; see Figure 1A). During the preparation phase of the TSST (T4), anticipatory cognitive appraisal (PASA) [38] was also assessed. Lastly, after completion of the TSST (T5), participants’ satisfaction towards their performance at the speech and math tasks was assessed on a VAS scale ranging from “not at all” (+1) to “very much” (+101).

### Physiological measures of stress

As physiological stress biomarkers, salivary cortisol (hypothalamic–pituitary–adrenal [HPA] axis activity), salivary alpha-amylase and heart rate (autonomic nervous system [ANS] activity) were assessed. Heart rate was recorded using a chest strap (Polar H10; Polar Electro Oy, Kempele, Finland). Data were collected over a 10 min period at baseline, during the TSST, and during the Social Reward task. Values in each time window were then averaged for statistical analyses. Saliva samples were collected via passive drool method using Salicaps (IBL, Hamburg, Germany) at 6 time points (T1, T2, T3, T5, T6, T7; see Figure 1A). Free cortisol concentration in saliva was determined by using commercial luminescence immunosorbent assay (LUM; IBL, Hamburg, Germany). Salivary alpha-amylase activity was measured using a kinetic colorimetric test and reagents obtained from DiaSys Diagnostic Systems (Holzheim, Germany). For heart rate, salivary cortisol and alpha-amylase analyses, outliers were defined as subjects with a baseline value (T1) 3 SDs over the mean baseline of the sample. This procedure led to the exclusion of two participants for cortisol (1 MORPH, 1 PLB) and of one participant for alpha-amylase (MORPH).

### Drug effects on cognitive functions and side-effects

To assess potential drug effects on cognitive functions, participants completed the Trail Making Test (TMT) [39] and the Digit Symbol Substitution Test (DSST) [40] 55 min after drug administration. Regarding subjective drug effects, participants filled out a self-report questionnaire assessing nausea, dry mouth and other 24 possible side-effects on a 4-point Likert scale (with the anchors 1 = “not at all” and 4 = “very much”) at baseline (T1), as well as 50 min (T2) and 160 min (T7) after drug administration (see Figure 1A).

### Statistical analyses

Statistical analyses were conducted in R [41]. Drug effects on subjective and physiological stress measures were analyzed using linear mixed effects models (LMM) with Drug (MORPH, PLB) and Time as fixed effects and by-subject random intercepts. Drug effects on ratings of wanting and liking, and force exerted were analyzed with LMMs including Drug (MORPH, PLB) and Reward Level (high, low, and very vow in the case of liking) as fixed effects, and by-subject random intercepts and slopes for Reward Level. For EMG data, LMMs for each muscle and task phase (anticipation, consumption) were fitted, including Drug (MORPH, PLB), trial-by-trial Wanting (for anticipation) or Liking (for consumption) and Epoch (Anticipation Pre-Effort and Post-Effort for anticipation, Delivery and Relax for consumption) as fixed effects, and by-subject random intercepts and slopes for Wanting/Liking, Epoch, and their interaction. In case of model unconvergence or singularity, random effects with the lowest cumulative variance were removed and, in case of categorical variables, transformed into the corresponding complex random intercept [42]. Group differences in age, BMI, and personality traits, as well as drug effects on executive functions, PASA, satisfaction for the TSST performance and side-effects were assessed using independent two-sided t-tests.

LMMs were computed using the *lmer* function of the lme4 package [43]. Type-III F-tests were computed with the Satterthwaite degrees of freedom approximation, using the *anova* function of the lmerTest package [44]. Significant interactions were further analyzed with multiple comparisons using the function *emmeans* from the homonymous package [45]. Results from all LMMs and multiple comparisons were controlled for the false discovery rate (FDR) associated with multiple testing using the Benjamini–Hochberg method [46].

The data and analysis scripts supporting the article are available at https://osf.io/gbd24/.

## Results

### Drug blinding

After completing the session, 50% of the participants who had actually received morphine correctly guessed to have received the drug. 55% of the total sample believed to have been administered with placebo, 29% with morphine, and 16% with naltrexone^2^. Overall, these numbers indicate successful blinding.

### Effects of morphine on cognitive functions and drug side-effects

There were no significant differences in the DSST and TMT scores across groups, indicating that drug administration did not have negative effects on attention, psychomotor and processing speed, and visuo-perceptual functions (See Table S1 in *Supplementary Material*). Morphine administration significantly increased self-reported weakness (MORPH vs. PLB at T2: *t*(69.9) = 2.64, *p* = 0.01, and at T7: *t*(55.96) = 2.19, *p =* 0.03) and dry mouth (MORPH vs. PLB at T7: *t*(57.70) = 2.56, *p* = 0.01). For all side-effect measures, group average scores were generally low, and no side-effect was on average rated as moderate or strong (see Figure S2 in *Supplementary Material*).

### Effects of morphine on stress response

#### Physiological measures

Morphine administration suppressed the cortisol response to the TSST (Drug*Time: *F*(5,375) = 7.68, FDR *p* < 0.001, Figure 2A). Specifically, the morphine group showed reduced salivary cortisol compared to the placebo group at T2 (FDR *p* = 0.079), T3 (FDR *p* < 0.01), T5 (FDR *p* < 0.001), T6 (FDR *p* < 0.001) and T7 (FDR *p* < 0.001) (Figure 2A). No drug effects were observed for salivary alpha-amylase (all *F* < 0.38, all FDR *p* > 0.62) or heart rate (all *F* < 2.83, all FDR *p* > 0.09).

**Figure 2.**
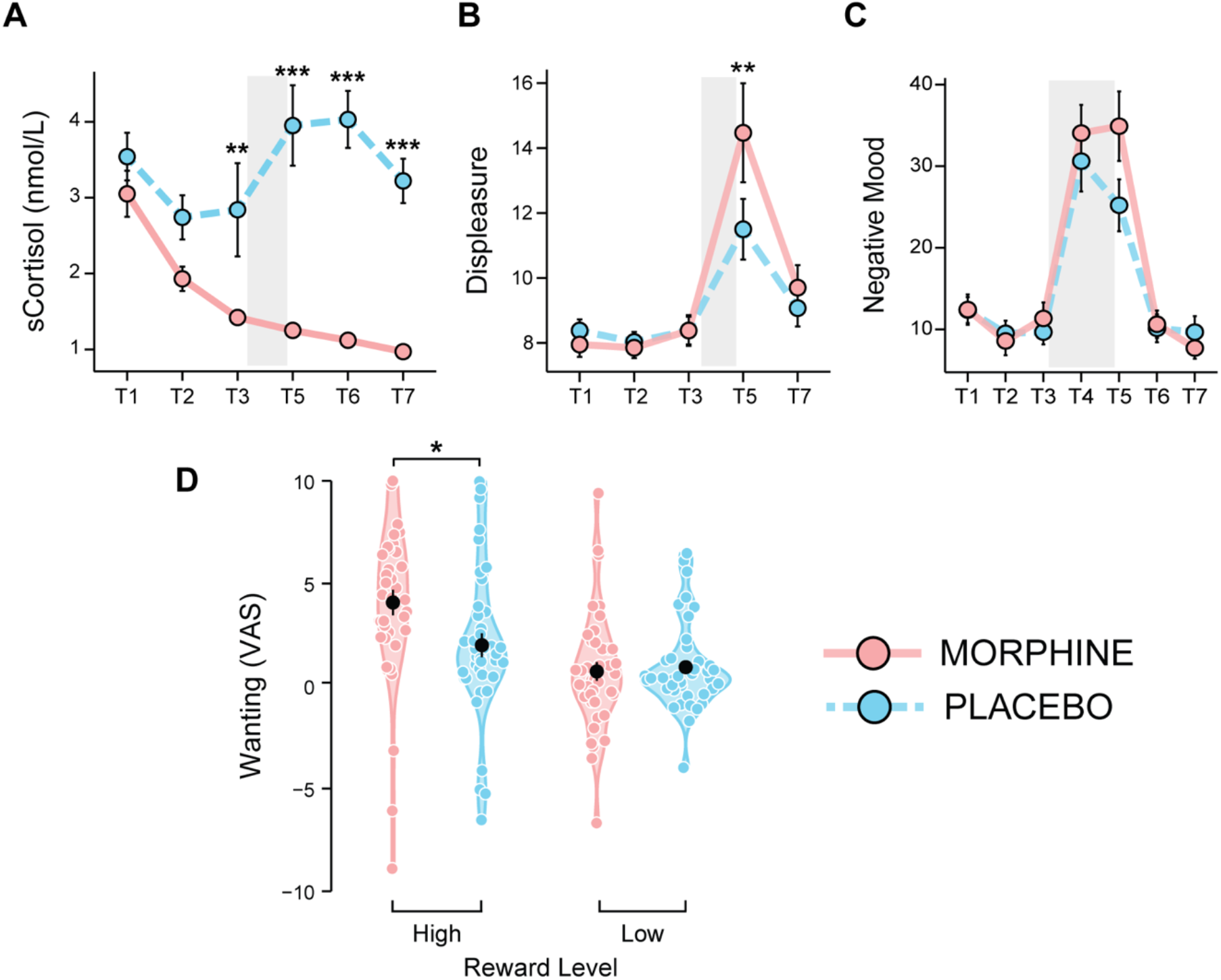
Effects of morphine administration on stress responses and social reward processing. (A) Morphine administration suppressed the hypothalamic–pituitary–adrenal (HPA) axis activity, as shown by blunted salivary cortisol (sCortisol) before and after stress induction. Morphine administration also increased the subjective negative response to stress, as shown by (B) higher scores on the “Displeasure” subscale (POMS) and (C) elevated negative mood (measured via visual analog scales) immediately after the Trier Social Stress Test (TSST). (D) Morphine administration resulted in significantly greater wanting for the high social reward (6 cm/s, CT-optimal touch), compared to placebo (black dots represent group means and colored dots represent individual means). Grey bars represent the TSST time window (anticipation, speech, and arithmetic task); error bars represent standard error of the mean; asterisks indicate significant differences between drug groups (* *p* <.05, *** *p* < .001).

#### Subjective measures

Morphine administration resulted in significantly higher scores of the POMS subscale “Displeasure” after stress induction (Drug*Time: *F*(4,312) = 2.97, FDR *p* = 0.03; MORPH vs. PLB at T5: FDR *p* < 0.01) (Figure 2B). A similar pattern, though not reaching the statistical significance threshold, was observed for negative mood (Drug*Time: *F*(6,468) = 2.16, FDR *p* = 0.068) (Figure 2C). No significant group differences were observed for positive mood or for the other POMS subscales (all FDR *p* > .15). Furthermore, no drug effects were observed in anticipatory stress (PASA primary and secondary appraisal at T4; both *t* < 0.60, *p* > 0.55), nor in the participants’ satisfaction towards their performance expressed after TSST completion (T5; *t*(75.2) = −0.63, *p* = 0.53).

#### Correlations between physiological and subjective measures of stress

Given the observed opposite effect of the drug on cortisol and mood, we conducted a correlation analysis to investigate the association between endocrine and subjective measures of stress. To this aim, we first computed the area under the curve in respect to increase (AUCi, from T3 to T7) [47] of the cortisol levels and of the negative mood ratings (VAS and POMS). Results showed a negative correlation between salivary cortisol and negative mood (VAS; *r*_s_ = −.36, *p bonferroni* = .048) as well as between cortisol and the Displeasure subscale of the POMS (*r*s = −.37, *p bonferroni* = .044), suggesting an inverse relationship between HPA axis and mood responses to stress.

### Effects of morphine on social reward processing

We first examined whether the number of high, low and very low rewards obtained during the task was similar for the two groups. A linear model on number of trials was fitted, including Drug (MORPH, PLB) and Reward Level (high, low, and very low). We observed only a main effect of Reward Level (*F*(2,369.5) = 11.28, *p* < 0.001), indicating that participants obtained more often high rewards as compared to low and very low rewards. We also tested for group differences in MVC, assessed before and after the Social Reward task and used to calibrate the effort task. No significant differences emerged (all *t* < −0.84, all *p* > 0.40), indicating no drug effects on participants’ grip force at rest. Last, we tested for group differences in the baseline activity of the corrugator and zygomaticus muscles. No significant differences emerged (all *t* < 1.27, all *p* > 0.21).

#### Effects of morphine on subjective ratings of wanting and liking

##### Ratings of Wanting

Participants administered morphine expressed significantly greater wanting of the high social reward (CT-optimal touch) compared to the placebo group (Drug*Reward Level: *F*(1,78) = 10.56, FDR *p* = 0.003; high social reward MORPH vs. PLB: FDR *p* = 0.035) (Figure 2D). No drug effects emerged for the low social reward (FDR *p* = 0.72).

##### Ratings of Liking

Participants in both groups expressed greater liking for high compared to low social rewards, which in turn were more liked than the very low social rewards (Reward Level: *F*(2,72.9) = 20.93, FDR *p* < 0.001; high vs. low vs. very low: all FDR *p* < 0.001). No significant effects of drug were observed (all *F* < 4.23, all FDR *p* > 0.07).

#### Effects of morphine on force exerted and hedonic facial reactions

We further assessed wanting of social rewards in terms of force exerted to obtain the tactile stimuli, as well as hedonic facial reactions during reward anticipation (Anticipation Pre- and Post-Effort phases). Hedonic facial reactions during and after consumption (Delivery and Relax phases) of the social reward were employed as a measure of liking.

##### Force exerted

Participants overall exerted greater force to obtain the high compared to the low social reward (Reward Level: *F*(1,78) = 14.23, FDR *p* < 0.001). No significant drug effects were observed (all *F* < 4.13, all FDR *p* > 0.07).

##### Facial EMG

No significant drug effects were observed on the activity of the corrugator and zygomaticus muscles during reward anticipation and consumption. The analyses on the zygomaticus revealed a significant Wanting by Epoch interaction (*F*(1,3831) = 10.61, FDR *p* < 0.01), as smiling was positively associated to the ratings of wanting during the announcement of the attained reward (Anticipation Pre-Effort) but not during the announcement of the attainable reward (Anticipation Pre-Effort). The analyses on the corrugator revealed a significant main effect of Epoch (*F*(1,73.1) = 8.79, FDR *p* = 0.03), as participants frowned less during the delivery of social touch as compared to the relax period.

## Discussion

Social motivation is a powerful force guiding behavior, as social rewards (e.g., social contact, bonding, affiliation) are fundamental to the individual’s physical and psychological well-being. Despite the important role of social contact in stress resilience, the neurochemical mechanisms underlying social contact seeking following stress exposure in humans are still poorly understood. In this study, we pharmacologically challenged the μ-opioid receptor (MOR) system to investigate its role in the regulation of the motivational and hedonic components of social reward processing following stress induction. To parallel previous animal research, participants were exposed to a stressor of social nature and interpersonal touch was used as a social reward. Further, force exerted to obtain the reward as well as hedonic facial reactions during its anticipation and consumption were assessed, together with subjective ratings of wanting and liking. Following the enhancement of the MOR system activity via administration of its agonist morphine, we observed suppression of the HPA axis activity (as indicated by a reduced cortisol response) and increased negative affect in response to psychosocial stress. Notably, this increased negative response to stress after morphine administration was followed by enhanced motivation for the most pleasurable social reward.

### Morphine-blunted cortisol stress response is associated with increased negative affect

Administration of the μ-opioid agonist morphine prior to TSST exposure resulted in blunted salivary cortisol response, indicating suppression of the HPA axis reactivity to stress. This is in line with previous animal and human evidence indicating an inhibitory role of μ-opioids on cortisol release following stress exposure [48]. Recently, two studies investigated the effects of partial (buprenorphine) and full (hydromorphone) MOR agonists on psychosocial stress, induced via TSST [14,15]. Akin to the present findings, reduced cortisol responses to stress were observed.

Buprenorphine and hydromorphone also reduced the perceived threat and the appraisal of how challenging the participants found the stress task, respectively. While the authors interpret the findings on cortisol and stress appraisal as indicators of a reduced stress response, no mood-buffering effects of the drug were observed. This is in contrast with the current study, where we find that the dampened cortisol response is accompanied by an enhanced aversive reaction to stress, as shown by higher negative affect following morphine compared to placebo administration. Human cortisol permeates the blood-brain barrier to feedback the central nervous system, reducing HPA axis activity and promoting emotion regulation [49,50]. The observed inverse relationship between the HPA axis activity and negative affect possibly indicates a disruption of this feedback loop by morphine administration and is in line with a mood-buffering effect of cortisol [49,51]. Accordingly, it was recently shown that pharmacological HPA axis suppression, by means of dexamethasone administration, blunts the cortisol response to stress and increases negative mood, especially in women [52,53].

Our results are opposite to our *a priori* hypothesis based on previous evidence from animal studies on separation distress indicating a reduction of stress indices, such as distress vocalizations, following MOR agonist administration [54]. The paradoxical morphine effect of enhanced aversive stress reaction observed in the current study vs. the previous animal literature may be explained by experimental differences, such as route and timing of administration of the opioid compounds. For instance, in animal research morphine is typically delivered intravenously after stress induction, possibly allowing the system to prepare to face the stressor via an initial mounting of the physiological stress response. In this study, morphine was administered orally to minimize the invasiveness of the administration procedure. Unlike intravenous administration, per-oral morphine has a slow pharmacokinetic profile and requires an average time of 60 min to reach the peak blood concentration. As the subjective response to acute stress, especially its effect on mood, typically lasts only for short periods of time after laboratory stress induction, orally administering the drug and waiting for it to reach peak concentration after the TSST would not have been feasible. For these reasons we administered the compound prior to stress exposure. However, the resulting suppression of the HPA axis activity before the beginning of the stress induction may have led to the observed difference in the mood response. Accordingly, the raising of cortisol levels, via cortisol administration, prior to stress induction has been shown to have a protective role in women, lowering the negative subjective reaction to stress [51].

### Morphine-induced increased aversive reaction to stress is associated with enhanced social motivation

Regarding the social reward task, the observed effects are consistent with previous models [3,4] suggesting that, under distress conditions, individuals seek physical social contact to down-regulate the negative state and re-store comfort. Accordingly, we showed that the morphine-induced increased negative reaction to the TSST was accompanied by enhanced social contact seeking. Specifically, we observed greater subjective wanting of the most pleasurable social reward (CT-optimal touch) following morphine administration, compared to placebo. Physical social contact, such as grooming in animals and caressing in humans, has a soothing function and is considered a powerful means to buffer distress and re-store comfort [55]. Interpersonal touch, and particularly slow CT-optimal touch in humans, has been shown to reduce the physiological signs of distress [e.g., 56,57] and the psychological consequences of aversive social situations, such as social rejection [58], representing therefore an especially appetitive stimulus in such situations. However, while previous models predict that the relief of negative affect through social contact is mediated by μ-opioids [3,4], due to the employed design and the opposite effects of the drug on stress responses compared to previous animal research, the role of the MOR system in the observed enhancement of contact seeking remains to be clarified.

The results are also consistent with existent evidence on the effects of stress and negative affect on reward processing. Previous research has indeed shown that stress exposure, and the consequent increase of negative affect, boosts the incentive value of appetitive stimuli, such as food, money, or drug cues. This results in a selective increased motivation to obtain the reward, despite an absent increase in the pleasure experienced once it is consumed [e.g., 17,59,60]. Notably, the results reported here replicate previous findings from our group indicating that, as already shown for non-social rewards, psychosocial stress increases wanting, but not liking, of rewards of social nature, such as interpersonal touch [16]. As in Massaccesi et al. 2021, the increased negative mood was accompanied by an increase in the subjective desire of receiving pleasurable social touch, but not in the force exerted to obtain it. While implicit wanting can occur without a conscious experience and mostly depends on dopaminergic mechanisms, explicit (subjective) wanting, or cognitive desire, requires awareness and is strongly linked to the individual’s previous liking experiences with and memories of the rewarding stimulus [61]. The cognitive desire of the stimulus is therefore driven by the pleasure that individuals expect to receive once the reward is consumed, based on these memories. Accordingly, a recent study on smokers showed that self-reported wanting and anticipated liking in response to smoking-related cues were strongly correlated and increased in heavy smokers, while this was not the case for consummatory liking, i.e. the pleasure reported after smoking a cigarette [62].

As expected, the effect of enhanced wanting for the social reward was selective for the most pleasurable touch (high reward level), characterized by slow stroking velocity (CT-optimal touch). This is different from our previous work [16], were a general increase of wanting, independent of reward magnitude, was observed following stress. The potentiation of wanting for the most pleasurable touch observed here may be due to a specific influence of the enhanced MOR system activity. Indeed, previous research has shown that pharmacological challenges of the MOR system have strongest effects on rewards of greatest magnitude [e.g., 6,18]. Therefore, it can be hypothesized that while stress alone may result in increased motivation for all available rewarding stimuli, an additional potentiation of motivation for the most valuable stimulus is seen for the best reward due to MOR stimulation. Alternatively, a reason for the partially different result pattern might be that, differently from the current study, in Massaccesi et al. 2021 participants were allowed to individually rank the three types of touch as high, low and very low. The high reward was therefore not necessarily the CT-optimal touch. This could have masked a possible impact of the type of touch. While our previous work included different state manipulations, this study focused on the effects of stress. Since slow CT-optimal touch is more likely to convey affiliative intentions such as social support compared to faster touch [63], here the slow CT-optimal touch was kept as fixed high reward to avoid possible confounding effects related to the speed of stroking.

### Study limitations

Some limitations of the study should be considered. First, while the use of a within-subject design is usually preferable in pharmacology, in this study we employed a between-subjects design. The choice was mainly motivated by the fact that repeated exposure to the TSST, especially with a short time interval, can lead to habituation of the stress response as well as other repetition effects, resulting in low test re-test reliability [e.g., 64,65] and thus outweighing by far the possibly higher statistical power a within-subject design might have had in principle. Second, despite our efforts in enhancing the social properties of the administered touch (e.g., skin-to-skin administration), the absence of a social relationship between the toucher and the participant, as well as other methodological aspects (e.g., use of a curtain separating participant and toucher), might have affected our results. Nevertheless, in humans an involvement of the MOR system in bond formation, rather than just maintenance, has also been observed [12]. Third, the current study tested a sample of healthy female participants, preventing a generalization to male individuals. Finally, it is unlikely that a complex behavior such as social motivation can be fully explained by the activity of a single neurochemical system only. For instance, the neuropeptides oxytocin and vasopressin are well-known for their crucial role in bonding and reproductive functions [66]. Stress exposure induces a potent activation of the dopaminergic system [67,68], and it has been shown that MOR stimulation disinhibits dopaminergic neurons in the ventral tegmental area and increases dopamine release in the nucleus accumbens [69]. To investigate how these systems interact in driving social contact seeking during aversive states seems therefore necessary to understand the neurobiology of social motivation.

### Conclusions

To conclude, our findings show that enhancing the MOR system activity before psychosocial stress exposure increases, rather than reduces, the aversive reaction to stress, leading to an increased subjective motivation for the attainment of highly rewarding social contact. Specifically, we showed that morphine administration blunted cortisol reactivity to stress and increased the negativity of the adverse experience, in line with a mood-buffering effect of cortisol. Further, the results indicate that this enhanced morphine-induced stress response was followed by increased desire to attain social rewards.

Overall, the results significantly extend previous human evidence on the modulation of the physiological and subjective responses to social stress by the MOR system and indicate a state-dependent regulation of social motivation. A better understanding of the effects of opioids on mood, wellbeing and social behaviors via opioid manipulations in healthy, opioid-naïve individuals may have important implications as drugs that act on the opioid system are widely consumed in both medical and non-medical contexts. To clarify the role of the MOR system in modulating contact seeking behaviors under distress, future studies should investigate the effects of MOR agonists and antagonists administered after stress induction (for instance via intravenous administration), as well as the interaction with other neurochemical systems, such as dopamine.

## Acknowledgments

We thank Gheorghe L. Preda and Carina Bum for their contribution in carrying out the medical procedures during data collection, Sebastian Korb, Emilio Chiappini, and Nace Mikus for valuable intellectual input, Nadine Skoluda for her support with the saliva analysis, and all the students who contributed to data collection.

## Authors contribution

CM: Conceptualization of the study, Data curation, Software, Formal analysis, Supervision, Investigation, Visualization, Methodology, Project administration, Funding acquisition, Writing - original draft, review and editing; MW: Contribution to the conceptualization of the design, Project administration, Medical testing, Funding acquisition, Writing - review and editing; BBQ: Contribution to the conceptualization of the design, Writing - review and editing; UMN: Contribution to the conceptualization of the design, Curation of saliva analysis, Writing - review and editing; CL: Contribution to the conceptualization of the design, Writing - review and editing; DM: Curation of blood analyses; GS: Conceptualization of the study, Supervision, Methodology, Funding acquisition, Project administration, Writing - review and editing.

All authors approved the final version of the manuscript.

## Footnotes

1 Due to technical problems, EMG data from 6 (4 MORPH) participants and HR data from 1 participant (MORPH) were not recorded. Saliva samples from one participant (MORPH) are also missing.

2 To reduce drug-related expectancy, participants were told they might receive an opioid agonist (morphine), antagonist (naltrexone) or placebo (but in reality could receive only morphine or placebo).

## References

1. Cacioppo JT, Cacioppo S. Social Relationships and Health: The Toxic Effects of Perceived Social Isolation. Soc Personal Psychol Compass. 2014;8:58–72.

2. Uchino BN. Social Support and Health: A Review of Physiological Processes Potentially Underlying Links to Disease Outcomes. J Behav Med. 2006;29:377–387.

3. Panksepp J. Affective Neuroscience: The Foundations of Human and Animal Emotions. Oxford University Press; 1998.

4. Loseth GE, Ellingsen D-M, Leknes S. State-dependent μ-opioid modulation of social motivation. Front Behav Neurosci. 2014;8:430.

5. Buchel C, Miedl S, Sprenger C. Hedonic processing in humans is mediated by an opioidergic mechanism in a mesocorticolimbic system. ELife. 2018;7:e39648.

6. Chelnokova O, Laeng B, Eikemo M, Riegels J, Løseth G, Maurud H, et al. Rewards of beauty: the opioid system mediates social motivation in humans. Mol Psychiatry. 2014;19:746–747.

7. Korb S, Götzendorfer SJ, Massaccesi C, Sezen P, Graf I, Willeit M, et al. Dopaminergic and opioidergic regulation during anticipation and consumption of social and nonsocial rewards. ELife. 2020;9:e55797.

8. Løseth GE, Eikemo M, Leknes S. Effects of opioid receptor stimulation and blockade on touch pleasantness: a double-blind randomised trial. Soc Cogn Affect Neurosci. 2019;14:411–422.

9. Inagaki TK, Irwin MR, Eisenberger NI. Blocking opioids attenuates physical warmth-induced feelings of social connection. Emot Wash DC. 2015;15:494–500.

10. Inagaki TK, Ray LA, Irwin MR, Way BM, Eisenberger NI. Opioids and social bonding: naltrexone reduces feelings of social connection. Soc Cogn Affect Neurosci. 2016;11:728–735.

11. Inagaki TK, Hazlett LI, Andreescu C. Opioids and social bonding: Effect of naltrexone on feelings of social connection and ventral striatum activity to close others. J Exp Psychol Gen. 2020;149:732–745.

12. Tchalova K, MacDonald G. Opioid receptor blockade inhibits self-disclosure during a closeness-building social interaction. Psychoneuroendocrinology. 2020;113:104559.

13. Hsu DT, Sanford BJ, Meyers KK, Love TM, Hazlett KE, Wang H, et al. Response of the μ-opioid system to social rejection and acceptance. Mol Psychiatry. 2013;18:1211–1217.

14. Bershad AK, Miller MA, Norman GJ, de Wit H. Effects of opioid- and non-opioid analgesics on responses to psychosocial stress in humans. Horm Behav. 2018;102:41–47.

15. Bershad AK, Seiden JA, de Wit H. Effects of buprenorphine on responses to social stimuli in healthy adults. Psychoneuroendocrinology. 2016;63:43–49.

16. Massaccesi C, Korb S, Skoluda N, Nater UM, Silani G. Effects of Appetitive and Aversive Motivational States on Wanting and Liking of Interpersonal Touch. Neuroscience. 2021;464:12–25.

17. Pool E, Brosch T, Delplanque S, Sander D. Stress increases cue-triggered ‘wanting’ for sweet reward in humans. J Exp Psychol Anim Learn Cogn. 2015;41:128–136.

18. Eikemo M, Løseth GE, Johnstone T, Gjerstad J, Willoch F, Le+knes S. Sweet taste pleasantness is modulated by morphine and naltrexone. Psychopharmacology (Berl). 2016;233:3711–3723.

19. Morrison I. Keep Calm and Cuddle on: Social Touch as a Stress Buffer. Adapt Hum Behav Physiol. 2016;2:344–362.

20. Fillingim RB, Gear RW. Sex differences in opioid analgesia: clinical and experimental findings. Eur J Pain. 2004;8:413–425.

21. Zubieta J-K, Dannals RF, Frost JJ. Gender and Age Influences on Human Brain Mu-Opioid Receptor Binding Measured by PET. Am J Psychiatry. 1999;156:842–848.

22. Kelly MM, Tyrka AR, Anderson GM, Price LH, Carpenter LL. Sex differences in emotional and physiological responses to the Trier Social Stress Test. J Behav Ther Exp Psychiatry. 2008;39:87–98.

23. Stier DS, Hall JA. Gender differences in touch: An empirical and theoretical review. J Pers Soc Psychol. 1984;47:440–459.

24. Suvilehto JT, Glerean E, Dunbar RIM, Hari R, Nummenmaa L. Topography of social touching depends on emotional bonds between humans. Proc Natl Acad Sci. 2015;112:13811–13816.

25. Freitag CM, Retz-Junginger P, Retz W, Seitz C, Palmason H, Meyer J, et al. Evaluation der deutschen Version des Autismus-Spektrum-Quotienten (AQ) - die Kurzversion AQ-k. Z Für Klin Psychol Psychother. 2007;36:280–289.

26. Spielberger CD, Gorsuch RL, Lushene RE. Manual for the State-Trait Anxiety Inventory. 1970. 1970.

27. Heimberg RG, Horner KJ, Juster HR, Safren SA, Brown EJ, Schneier FR, et al. Psychometric properties of the Liebowitz Social Anxiety Scale. Psychol Med. 1999;29:199–212.

28. Wilhelm FH, Kochar AS, Roth WT, Gross JJ. Social anxiety and response to touch: incongruence between self-evaluative and physiological reactions. Biol Psychol. 2001;58:181–202.

29. World Medical Association. World Medical Association Declaration of Helsinki: ethical principles for medical research involving human subjects. JAMA. 2013;310:2191–2194.

30. Sheehan DV, Lecrubier Y, Sheehan KH, Amorim P, Janavs J, Weiller E, et al. The Mini-International Neuropsychiatric Interview (M.I.N.I.): The Development and Validation of a Structured Diagnostic Psychiatric Interview for DSM-IV and ICD-10. J Clin Psychiatry. 1998;59:0–0.

31. Labuschagne I, Grace C, Rendell P, Terrett G, Heinrichs M. An introductory guide to conducting the Trier Social Stress Test. Neurosci Biobehav Rev. 2019;107:686–695.

32. Lugo RA, Kern SE. Clinical Pharmacokinetics of Morphine. J Pain Palliat Care Pharmacother. 2002;16:5–18.

33. Kirschbaum C, Pirke KM, Hellhammer DH. The ‘Trier Social Stress Test’--a tool for investigating psychobiological stress responses in a laboratory setting. Neuropsychobiology. 1993;28:76–81.

34. Löken LS, Wessberg J, Morrison I, McGlone F, Olausson H. Coding of pleasant touch by unmyelinated afferents in humans. Nat Neurosci. 2009;12:547–548.

35. Korb S, Massaccesi C, Gartus A, Lundström JN, Rumiati R, Eisenegger C, et al. Facial responses of adult humans during the anticipation and consumption of touch and food rewards. Cognition. 2020;194:104044.

36. Fridlund AJ, Cacioppo JT. Guidelines for Human Electromyographic Research. Psychophysiology. 1986;23:567–589.

37. Albani C, Blaser G, Geyer M, Schmutzer G, Brähler E, Bailer H, et al. The German short version of ‘Profile of Mood States’ (POMS): psychometric evaluation in a representative sample. PPmP - Psychother · Psychosom · Med Psychol. 2005;55:324–330.

38. Gaab J, Rohleder N, Nater UM, Ehlert U. Psychological determinants of the cortisol stress response: the role of anticipatory cognitive appraisal. Psychoneuroendocrinology. 2005;30:599–610.

39. Reitan RM. Validity of the Trail Making Test as an Indicator of Organic Brain Damage. Percept Mot Skills. 1958;8:271–276.

40. Wechsler D. The measurement of adult intelligence. Baltimore, MD, US: Williams & Wilkins Co; 1939.

41. R Core Team. R: A language and environment for statistical computing. R Foundation for Statistical Computing; 2019.

42. Scandola M, Tidoni E. The development of a standard procedure for the optimal reliability-feasibility trade-off in Multilevel Linear Models analyses in Psychology and Neuroscience. 2021.

43. Bates D, Mächler M, Bolker B, Walker S. Fitting Linear Mixed-Effects Models Using lme4. J Stat Softw. 2015;67:1–48.

44. Kuznetsova A, Brockhoff PB, Christensen RHB. lmerTest Package: Tests in Linear Mixed Effects Models. J Stat Softw. 2017;82:1–26.

45. Lenth RV. emmeans: Estimated marginal means, aka least-squares means. R package version 1.5. 4. 2021.

46. Benjamini Y, Hochberg Y. Controlling the False Discovery Rate: A Practical and Powerful Approach to Multiple Testing. J R Stat Soc Ser B Methodol. 1995;57:289–300.

47. Pruessner JC, Kirschbaum C, Meinlschmid G, Hellhammer DH. Two formulas for computation of the area under the curve represent measures of total hormone concentration versus time-dependent change. Psychoneuroendocrinology. 2003;28:916–931.

48. Pechnick RN. Effects of Opioids on the Hypothalamo-Pituitary-Adrenal Axis. Annu Rev Pharmacol Toxicol. 1993;33:353–382.

49. Het S, Schoofs D, Rohleder N, Wolf OT. Stress-Induced Cortisol Level Elevations Are Associated With Reduced Negative Affect After Stress: Indications for a Mood-Buffering Cortisol Effect. Psychosom Med. 2012;74:23–32.

50. McEwen BS. Stress, adaptation, and disease. Allostasis and allostatic load. Ann N Y Acad Sci. 1998;840:33–44.

51. Het S, Wolf OT. Mood changes in response to psychosocial stress in healthy young women: effects of pretreatment with cortisol. Behav Neurosci. 2007;121:11–20.

52. Ali N, Nitschke JP, Cooperman C, Pruessner JC. Suppressing the endocrine and autonomic stress systems does not impact the emotional stress experience after psychosocial stress. Psychoneuroendocrinology. 2017;78:125–130.

53. Ali N, Cooperman C, Nitschke JP, Baldwin MW, Pruessner JC. The effects of suppressing the biological stress systems on social threat-assessment following acute stress. Psychopharmacology (Berl). 2020;237:3047–3056.

54. Panksepp J, Herman BH, Vilberg T, Bishop P, DeEskinazi FG. Endogenous opioids and social behavior. Neurosci Biobehav Rev. 1980;4:473–487.

55. Dunbar RIM. The social role of touch in humans and primates: Behavioural function and neurobiological mechanisms. Neurosci Biobehav Rev. 2010;34:260–268.

56. Pawling R, Cannon PR, McGlone FP, Walker SC. C-tactile afferent stimulating touch carries a positive affective value. PLOS ONE. 2017;12:e0173457.

57. Triscoli C, Croy I, Steudte-Schmiedgen S, Olausson H, Sailer U. Heart rate variability is enhanced by long-lasting pleasant touch at CT-optimized velocity. Biol Psychol. 2017;128:71–81.

58. von Mohr M, Krahé C, Beck B, Fotopoulou A. The social buffering of pain by affective touch: a laser-evoked potential study in romantic couples. Soc Cogn Affect Neurosci. 2018;13:1121–1130.

59. Kumar P, Berghorst LH, Nickerson LD, Dutra SJ, Goer FK, Greve DN, et al. Differential effects of acute stress on anticipatory and consummatory phases of reward processing. Neuroscience. 2014;266:1–12.

60. Lewis AH, Porcelli AJ, Delgado MR. The effects of acute stress exposure on striatal activity during Pavlovian conditioning with monetary gains and losses. Front Behav Neurosci. 2014;8.

61. Pool E, Sennwald V, Delplanque S, Brosch T, Sander D. Measuring wanting and liking from animals to humans: A systematic review. Neurosci Biobehav Rev. 2016;63:124–142.

62. Selby DL, Harrison AA, Fozard TE, Kolokotroni KZ. Dissociating wanting and anticipated liking from consummatory liking in smokers with different levels of nicotine dependence. Addict Behav. 2020;102:106185.

63. Kirsch LP, Krahé C, Blom N, Crucianelli L, Moro V, Jenkinson PM, et al. Reading the Mind in the Touch: Neurophysiological Specificity in the Communication of Emotions by Touch. Neuropsychologia. 2017. 29 May 2017. https://doi.org/10.1016/j.neuropsychologia.2017.05.024.

64. Gerra G, Zaimovic A, Mascetti GG, Gardini S, Zambelli U, Timpano M, et al. Neuroendocrine responses to experimentally-induced psychological stress in healthy humans. Psychoneuroendocrinology. 2001;26:91–107.

65. Petrowski K, Wintermann G-B, Siepmann M. Cortisol response to repeated psychosocial stress. Appl Psychophysiol Biofeedback. 2012;37:103–107.

66. Donaldson ZR, Young LJ. Oxytocin, Vasopressin, and the Neurogenetics of Sociality. Science. 2008;322:900–904.

67. Bloomfield MA, McCutcheon RA, Kempton M, Freeman TP, Howes O. The effects of psychosocial stress on dopaminergic function and the acute stress response. ELife. 2019;8:e46797.

68. Holly EN, Miczek KA. Ventral tegmental area dopamine revisited: effects of acute and repeated stress. Psychopharmacology (Berl). 2016;233:163–186.

69. Pan ZZ. μ-Opposing actions of the κ-opioid receptor. Trends Pharmacol Sci. 1998;19:94–98.

