## Supplementary Material for "Opioid-blunted cortisol response to stress is associated to increased negative mood and wanting of social reward"

**Index**

1. *Effects of COVID-19 pandemic*
   a. Safety measures
   b. Additional analyses to assess the effects of the COVID-19 pandemic and employed safety measures on responses to social rewards
2. *Drug effects on cognitive functions*
3. *Drug side-effects*
4. *Serum levels of morphine and its metabolites*

### 1. Effects of COVID-19 pandemic

Half of the sample was collected after the COVID-19 pandemic outbreak. The implemented safety measures, as well as the statistical analyses conducted to assess the possible effects of the pandemic on the study dependent variables are described below.

#### a. Safety measures

After the COVID-19 pandemic outbreak the following safety measures were implemented:

- All experimenters wore face masks. The evaluating panel of the TSST wore a special face mask with a clear plastic insert on the mouth region in order to allow the participant to see the "absence of facial feedback" (crucial for stress induction) during the stress paradigm.
- Participants were provided with clear mouth visors resting on the chin in order to minimize the contact with the face and avoid disturbances during facial electromyography (EMG).

In order to assess participants reaction towards these safety measures, they were asked to answer the following questions at the end of the experimental session:

1) "How well/comfortable did you feel during the study?" rated on a VAS ranging from 1 (not all) to 101 (very much).
2) "How high do you rate the risk of infection during the study?" rated on a VAS ranging from 1 (very high) to 101 (very low).
3) "Were you afraid of being infected with COVID-19 during the study?", Yes/No
4) "How comfortable did you feel with the research team wearing a mask during the study?" rated on a VAS ranging from 1 (not all) to 101 (very much).

As shown in Figure S1, participants did not report to feel threatened by COVID-19 infection during the study, and to feel comfortable during the session.

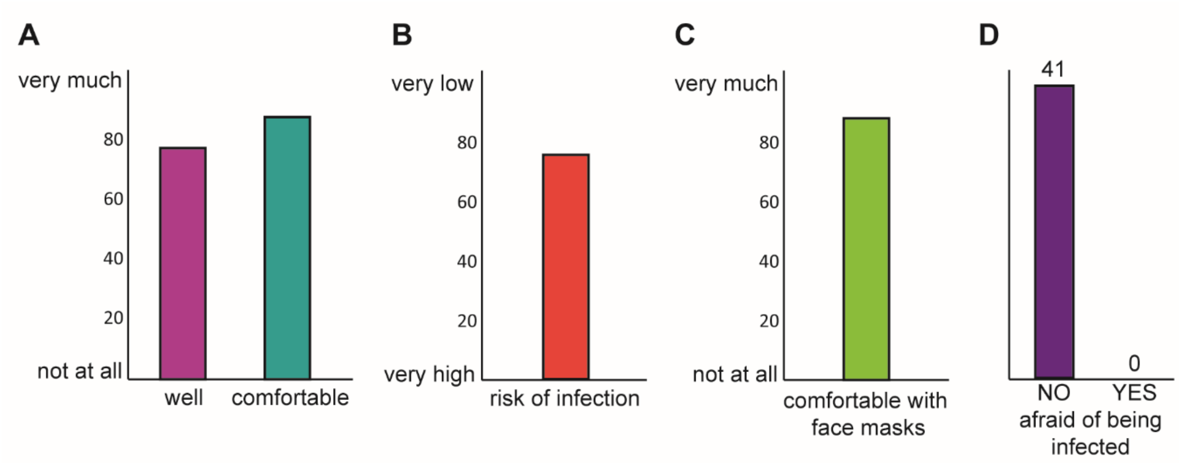


*Figure S1.* Participants' attitudes toward COVID-19 pandemic and safety measures during the study. (A) Ratings of wellness and comfort. (B) Perceived risk of COVID-19 infection during the study. (C) Ratings of comfort related to use of face masks during the study. (D) Perceived fear of having been infected with COVID-19 during the study.

#### b. Additional analyses to assess the effects of the COVID-19 pandemic and employed safety measures on responses to social rewards

We tested for possible effects of the pandemic and related implemented safety measures on the subjective stress response (positive and negative mood, POMS subscales), as well as on the ratings of wanting and liking, on the force exerted to obtain the social rewards, and on facial EMG data, by adding the covariate "COVID-19" (2 levels: pre-covid19, covid19) to the statistical models. No changes in the pattern of results were observed.

### 2. Drug effects on cognitive functions

*Table S1.* Mean (SD) scores at the Trial Making Test (TMT) part A and part B, and at the Digit Symbol Substitution Test (DSST) across drug groups.

| | MORPHINE | PLACEBO | *p* value |
|---|---|---|---|
| TMT A | 26.33 (8.02) | 26.40 (8.60) | 0.97 |
| TMT B | 57.25 (33.05) | 58.93 (21.94) | 0.79 |
| DSST | 54.20 (8.99) | 54.10 (9.32) | 0.96 |

### 3. Drug side-effects

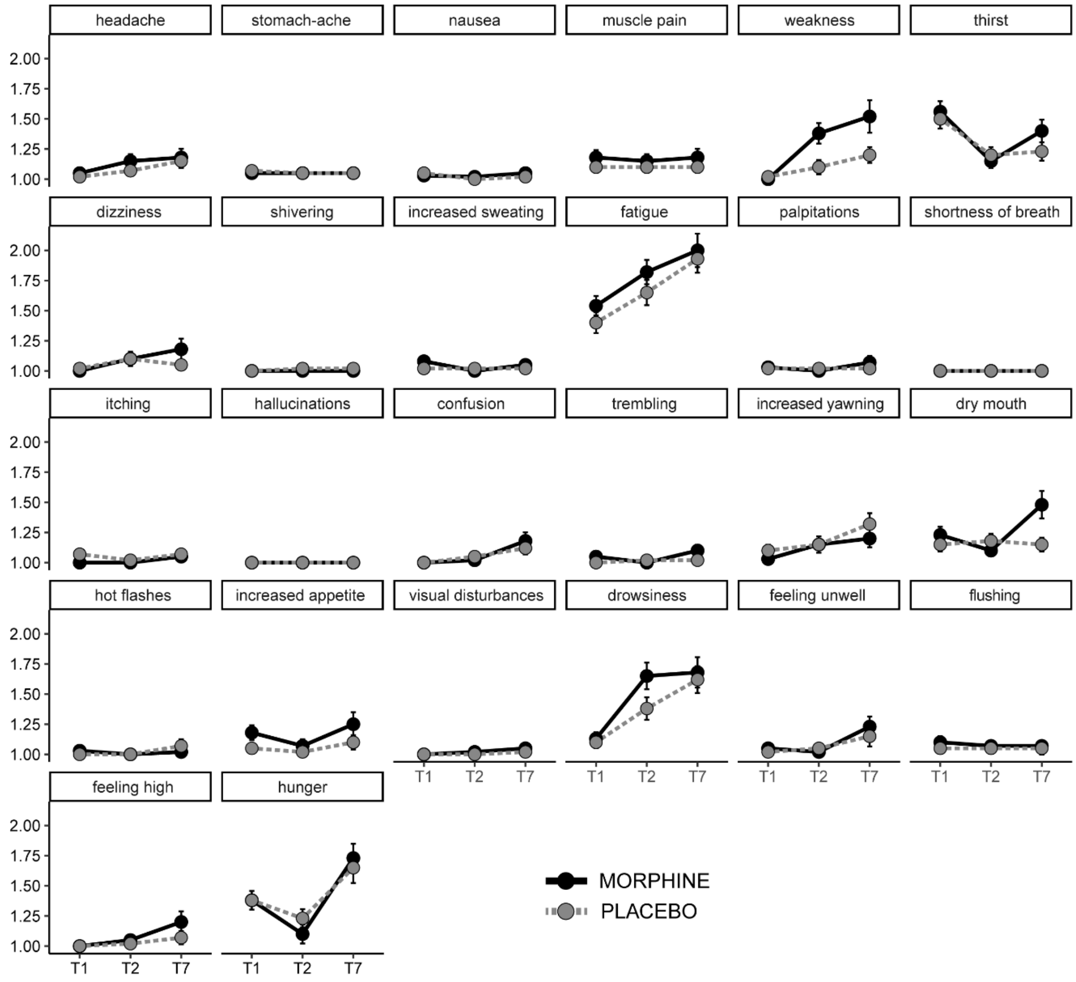


*Figure S2.* Drug side-effects assessed at baseline (T1), 60 min (T2) and 160 min (T7) after drug administration using a 4-point Likert scale (with the anchors 1 = "not at all" and 4 = "very much").

### 4. Serum levels of morphine and its metabolites

A blood sample was drawn at the end of the session (~180 min after drug administration). Analyses were performed at the Institute of Clinical Chemistry, University Hospital Zurich, using liquid chromatography coupled to mass spectrometry (LC-MS) to identify serum levels of morphine and its two major metabolites: morphine-3-glucuronide (M3G) and morphine-6-glucuronide (M6G). Blood samples from 3 participants could not be obtained and 10 samples were lost because of storage problems. Results from the available samples confirmed drug uptake, as shown in *Table S2*.

*Table S2.* Serum levels (nmol/l) of morphine and its major metabolites at the end of the experimental session (~180 min after drug administration). M3G & M6G = morphine-3/6-glucronide.

| | **M** | **SD** |
|---|---|---|
| Morphine | 12.63 | 6.82 |
| M3G | 394.79 | 179.46 |
| M6G | 84.63 | 44.32 |
